# Development of an orally-administrable tumor vasculature-targeting therapeutic using annexin A1-binding D-peptides

**DOI:** 10.1101/2020.10.12.335851

**Authors:** Motohiro Nonaka, Hideaki Mabashi-Asazuma, Donald L. Jarvis, Kazuhiko Yamasaki, Tomoya O. Akama, Masato Nagaoka, Toshio Sasai, Itsuko Kimura-Takagi, Yoichi Suwa, Takashi Yaegashi, Chun-Ten Huang, Chizuko Nishizawa-Harada, Michiko N. Fukuda

**Affiliations:** Laboratory for Drug Discovery, National Institute of Advanced Industrial Science and Technology, Tsukuba, Ibaraki 305-8568, Japan; Department of Biological Chemistry, Human Health Sciences, Graduate School of Medicine, Kyoto University, Kyoto, 606-8507, Japan; Department of Molecular Biology, University of Wyoming, Laramie, WY 82071, USA; Biomedical Research Institute, National Institute of Advanced Industrial Science and Technology, Tsukuba, Ibaraki 305-8566, Japan; Department of Pharmacology, Kansai Medical University, Hirakata, Osaka 573-1010, Japan; Yakult Central Institute, Kunitachi, Tokyo 186-8650, Japan; Cancer Center, Sanford-Burnham-Prebys Medical Discovery Institute, La Jolla, CA 92037, USA

## Abstract

IF7 peptide, which binds to the annexin A1 (ANXA1) N-terminal domain, functions as a tumor vasculature-targeted drug delivery vehicle after intravenous injection. To enhance IF7 stability *in vivo*, we undertook mirror-image peptide phage display using a synthetic D-peptide representing the Anxa1 N-terminus as target. Peptide sequences were identified, synthesized as D-amino acids, and designated as dTIT7, which was shown to bind the ANXA1 N-terminus. Whole body imaging of mouse brain tumors modeled with near infrared fluorescent IRDye-conjugated dTIT7 showed fluorescent signals in brain and kidney. Furthermore, orally-administered geldanamycin (GA)-conjugated dTIT7 suppressed brain tumor growth. Ours is a proof-of-concept experiment showing that Anxa1-binding D-peptide could be developed as an orally-administrable, tumor vasculature-targeted therapeutic.

Role of each author: MN designed and performed experiments, analyzed data, and wrote the manuscript; HMA and DLJ produced recombinant ANXA1 protein; KY conducted NMR analysis and data analysis; TOA designed, performed and analyzed LC-MS/MS data; MN, TS, IKT, YS, and TY analyzed peptide-binding assays and performed in silico structural analysis; CTU produced lentivirus for luciferase expression; CNH performed peptide binding assays, tissue culture and animal experiments; and MNF supervised the project and wrote the manuscript.

## Introduction

It is widely accepted that vasculature surfaces are heterogenous and express varying tissue-specific receptors under different pathological conditions [1]. Targeted-drug delivery to a disease-specific receptor on the endothelial cell surface could enable high therapeutic efficacy with minimum side effects. In order to enable drug delivery through intravenous route, it is essential to identify specific vasculature surface markers. Oh *et al*., used subtractive proteomics analysis of malignant vs. normal vasculature to identify Annexin A1 (ANXA1) as highly specific surface marker of malignant tumor vasculature [2, 3]. Coincidentally we found a linear 7-mer peptide IFLLWQR (IF7) that binds the ANXA1 N-terminus [4-6]. Upon intravenous injection into tumor-bearing mice, a conjugate of IF7 with the anti-cancer drug geldanamycin (GA) suppressed growth of prostate, breast, melanoma and lung tumors, and IF7-conjugated SN-38 suppressed colon cancer growth in mice at low dose without side effects [5]. Moreover, intravenously-injected IF7 accumulated on the tumor endothelial cell surface, was endocytosed into vesicles, and crossed tumor endothelial cells by transcytosis [5]. Thus we hypothesized that an IF7-conjugated drug would overcome the blood-brain-barrier (BBB) to eradicate brain tumors. Indeed, intravenous injection of the IF7-conjugated anti-tumor agent SN-38 into model mice harboring brain tumors efficiently reduced the size of brain tumor at low dosage, which apparently invoke host immune reaction against brain tumor leading into complete remission of brain tumor [6].

We conjugated IF7 to SN-38 through an esterase-cleavable linker, allowing SN-38 to be freed from the peptide once it reached the tumor vasculature. IF7 peptide itself was also susceptible to proteases. These properties of IF7-SN38 compromises its stability *in vivo* [5].

Here, to construct a protease-resistant form of IF7 that retains ANXA1-binding activity, we undertook mirror-image phage library screening taking an advantage of the fact that IF7 binds to chemically synthesized ANXA1 N-terminal domain (1-15 residues plus additional cysteine at 16), designated as MC16 [6]. This phage library screening identified the peptide dTIT7, which represents an ANXA1-binding D-type peptide. We then conjugated it to geldanamycin (GA) through an uncleavable linker to generate GA-dTIT7. We present proof-of-concept data showing that orally-administered GA-dTIT7 suppresses brain tumor growth in mice.

## Materials and Methods

### Materials

Unless noted, peptides used here, including the D-type peptides dTIT7, dLRF7, dSPT7, dLKG7 and dLLS7, were synthesized by GenScript (Piscataway, NJ). D-MC16 and L-MC16 peptides, with human 15 N-terminal ANXA1 residues plus a cysteine residue at 16 position (MAMVSEFLKQAWFIEC) and L-MC16 mutants were synthesized by Bio-Synthesis (Lewisville, TX). *Iso*dTIT7, in which prolines contain ^13^C and ^15^N, were synthesized by Peptide Institute, Osaka, Japan.

### Mirror-image phage library screening

Library screening strategies [7, 8] were adapted to identify a peptide sequence binding the ANXA1 N-terminal domain. D-MC16 peptide (described above) was chemically synthesized as D-amino acids, dissolved in DMSO, and used to coat maleimide-activated plates (Corning) at 10 nmol/well at 4°C for 20 hours. After blocking with SuperBlock solution (Thermo), screening was performed using a T7 phage library comprised of fully random 7-mer peptides, provided by Dr. E. Ruoslahti, Sanford-Burnham-Prebys Medical Discovery Institute (SBP). The phage peptide sequence was determined using an Ion Torrent Next Generation sequencer (Thermo). Top-ranked sequences TITWPTM or dTIT7 and the next four high-ranking peptides were then chemically synthesized using D-amino acids.

### Cell line

PGK-Luc lentiviral vector was produced at the Virus Core Facility of SBP. Rat glioma C6 and mouse melanoma B16F1cells were infected with lentivirus harboring firefly luciferase, to produce C6-Luc and B16F1-Luc lines [5, 6]. Lines were cultured in Dulbecco’s-Modified Eagle + F2 medium supplemented with 10% fetal bovine serum and 100 units each/mL penicillin and streptomycin, at 37C in a humidified 5% CO2 incubator.

### Binding of biotinylated D-peptides to L-MC16

To assess dTIT7 binding, wells of Sulfhydryl-BIND Surface Maleimide plates (Corning) were coated 20 hours with wild type (WT) and mutant L-MC16 in water at 4°C. After washing with PBS containing 0.02% Tween 20 (PBST), wells were blocked 1 hour with 10% superblock (Thermo) in PBST at room temperature. Biotinylated dTIT7 (10 µg dissolved in 10% superblock in PBST (1 mL) was added to each well at 100 µl/well prepared as above and incubated 30 min at room temperature for. After three PBST washes, 100 μl streptavidin-peroxidase (0.2 μg/ml) in 10% superblock in PBST containing 2% bovine serum albumin was added to each well and incubated 30 min. After three PBST washes, 100 μL of the peroxidase substrate one-step-TMB (Thermo) was added and incubated until the color developed. The reaction was stopped by adding 100 μl 2N sulfuric acid, and absorbance at 450 nm monitored using an ELISA plate reader.

### In silico conformational analysis of the ANXA1 N-terminus and dTIT7 docking

ANXA1 coordinates were obtained from the protein data bank (PDB). Both 1HM6 and 1MCX structures were derived from pig Anxa1 (89.6% sequence identity to human ANXA1 (Accession: P04083)). For the protein-protein docking structure of ANXA1, N-terminal free ANXA1 was built using MOE (Molecular Operating Environment) software ver. 2010.10 (Chemical Computing group). To obtain the dimer structure of ANXA1 with a free N-terminus, we used ZDOCK ver. 3.0.1 [9], which uses a Fast Fourier Transform-based algorithm to analyze proteins as rigid bodies during docking, searches for all possible binding orientations of a ligand along the receptor protein surface and provides docking poses ranked by Zdock scores associated with shape complementarity, desolvation and electrostatic properties. Hydrogen atoms of ANXA1 dimers calculated from Zdock were minimized using the AMBER99 force field.

### Preparation of recombinant ANXA1 protein

Recombinant, full-length ANXA1 was expressed using the baculovirus expression system [10], as described [6, 10]. Briefly, baculoviruses were prepared by recombining BacPAK6Δchi/cath baculovirus DNA with pAcP(-)-based baculovirus transfer vectors, which encode the transgene controlled by the baculovirus *p6*.*9* promoter. Recombinant ANXA1 protein harbored an N-terminal honeybee melittin signal peptide followed by a His_8_-tag and the enterokinase recognition sequence, DDDDR. Proteins were purified from Sf9 culture supernatants harvested 42 hours after infection using HisPur Ni-NTA resin (Pierce). Untagged ANXA1 was isolated by His-tagged enterokinase (Genscript) treatment followed by Ni-affinity chromatography. Protein concentration was determined using the BCA protein assay kit (Pierce).

### NMR Measurements

NMR was measured in solutions consisted of 50 μM peptide(s), 10 mM d_11_-Tris-HCl (pH 7.5) (Isotec Inc., IL), 150 mM NaCl, 1 mM d_10_-dithiothreitol (DTT) (Isotec Inc., IL), 0.1 mM sodium 2,2-dimethyl-2-silapentane-5-sulfonate (DSS), and 5% D_2_O. NMR spectra were recorded at 298 K on a Bruker (Germany) Avance III-500 spectrometer (^1^H frequency: 500.13 MHz). Chemical shifts were referenced to the peak of internal DSS. Relaxation time *T*_2_ was measured using the Carr-Purcell-Meiboom-Gill sequence and analyzed with the Topspin 3.2 program (Bruker). Error levels were estimated by four repeated experiments.

### Vertebrate animal use

Mouse protocols adhered to the NIH Guide for the Care and Use of Laboratory Animals and were approved by Institutional Review Committees at National Institute of Advanced Industrial Science and Technology (AIST) and Kyoto University School of Medicine in Japan. Experiments of brain tumor model mouse were conducted when tumor size determined as photon number was between 1 x10^4 and 1×10^7. When brain tumor grew more than 1×10^7, the mouse was euthanized by placing the animal under saturated isoflurane gas (1∼2 mL isoflurane in 250 mL chamber) followed by cervical dislocation. No animal died before meeting the criteria for euthanasia.

### Generation of brain tumor model mice

C6-Luc cells (4.8 ×10^4^ in 4 μl PBS) were injected into C57BL/6 mouse brain striatum using a stereotaxic frame as described [11]. Seven days later, mice underwent imaging for luciferase-expressing tumors. To do so, 100μl luciferin (30 mg/ml PBS) was injected peritoneally, and then mice were anesthetized under isoflurane gas (20 ml/min) supplemented with oxygen (1 ml/min) and placed under a camera equipped with a Xenogen IVIS 200 imager at AIST animal facility. Photon numbers were measured for 1-10 sec or for 1 min.

### Near infra-red fluorescence whole body imaging

Each 7-mer D-peptide with an N-terminal cysteine was synthesized by GenScript (Piscataway, NJ). Peptides were conjugated with IRDye 800CW maleimide (Li-Cor) through the cysteine residue at room temperature for 2 hours according to the manufacturer’s instruction. After reverse-phase HPLC purification, the conjugate was dissolved in DMSO and 6% glucose to a final concentration of 0.2 µM. The C6-Luc brain tumor model mouse was generated in nude mice as described above. When photon number reached 5×10^4^, each IRDye-conjugated D-peptide (100 μl) was injected intravenously through the tail vein. Near infra-red fluorescence in the mouse was monitored at 15 min after injection using an IVIS system and daily over 6 days.

### Conjugation of dTIT7 with a geldanamycin analogue

Procedures of Mandler *et al*. [12] were modified as follows. Geldanamycin (GA, 100 mg) was dissolved in chloroform (18 mL). 1, 3-diaminopropane (APA, 50 μl, molar ratio x 3.3 eq to GA) was also dissolved in chloroform (2 mL). APA solution was added slowly to GA and reacted at ambient temperature under argon gas for 20 hours. Hexane (100 mL) was then added slowly to precipitate a purple product (17-APA-GA or 17-DMAG), which was filtered through a glass filter. The precipitate was solubilized in chloroform (30 mL) and conjugated immediately to *N*-maleimidobutyril oxysuccinimide ester (GMBS, 100 mg) dissolved in chloroform (10 mL) and left at ambient temperature for 60 min under argon gas. The mixture was then concentrated on a rotary evaporator and applied to silica gel for thin layer chromatography with a solvent system of chloroform: methanol (9:1, v/v). A purple band representing GMB-APA-GA was isolated and extracted from the gel with methanol. GMB-APA-GA was further purified by C18 reverse phase HPLC with an acetonitrile gradient from 40-80% in water containing 0.1% trifluoro acetic acid. HPLC-purified GMB-APA-GA was dissolved in methanol (10 mL), and C-dTIT7 peptide (equimolar to GMB-APA-GA) was also dissolved in methanol (10 mL). Both were mixed at ambient temperature for 20 hours under argon gas. The product GA-dTIT7 (1719.52 Da) was purified by HPLC. GA-dTIT7 structure was validated by MALDI TOF-MS. Control GA-C (893.44 Da), GA-conjugated with cysteine only, was similarly prepared.

### LC-MS/MS analysis of GA-dTIT7 in mouse serum

C57BL/6 mice (8 week-old females) (n=6) were fasted overnight and then placed under isoflurane gas and administered a single dose of GA-dTIT7 (1 mg) dissolved with 10% taurodeoxycholate in water (200 μL) via oral gavage. Blood (50 μL) was collected from the facial vein at 0 min (pre-dose), 30 min, 60 min, 90 min and 120 min after administration using a lancet and placed into sodium heparin for plasma preparation. Then 1 μL *iso*-GA-dTIT7 (1 mg/mL in dimethylsulfoxide) was added as an internal standard to 9 μL plasma. After addition of cold acetone (40 μL), each sample was centrifuged to remove precipitates and an aliquot of supernatant was injected into an LC-MS/MS spectrometer.

### Oral administration of GA-dTIT7 to brain tumor-bearing mice

When photon numbers of B16-Luc or C6-Luc brain tumors reached 5×10^4^, oral administration of GA-dTIT7 or control GA-C was initiated. GA-dTIT7 (1719.52 Da, 2.0 mg) or GA-C (893.44 Da, 1.0 mg) was dissolved in 10 μL DMSO and diluted with 200 μL 10% taurodeoxycholate in water and then orally administered using a gavage.

### Statistical analysis

Statistical analyses were performed using GraphPad Prism program. Data sets were compared using Student’s unpaired *t*-test (two-tailed). A *p* value ≤ 0.05 was considered significant.

## Results

### Identification of linear 7-mer D-peptides by a mirror-image phage display screen

We showed previously that IF7 binds the Anxa1 N-terminal domain and that a chemically synthesized peptide representing this domain (designated MC16) was sufficient for IF7 binding [5, 6]. Here, we undertook mirror-image phage library screening for a protease-resistant D-type version of IF7 using synthetic D-MC16 peptide as target (Fig. 1A). This procedure resulted in enrichment for several phage clones (Fig. 1BCD), many showing a TITWPTM motif based on deep sequencing (Supplemental Table 1). We designated TITWPTM as TIT7 and a synthetic peptide of TIT7 composed of D-amino acids as dTIT7.

**Fig. 1.**
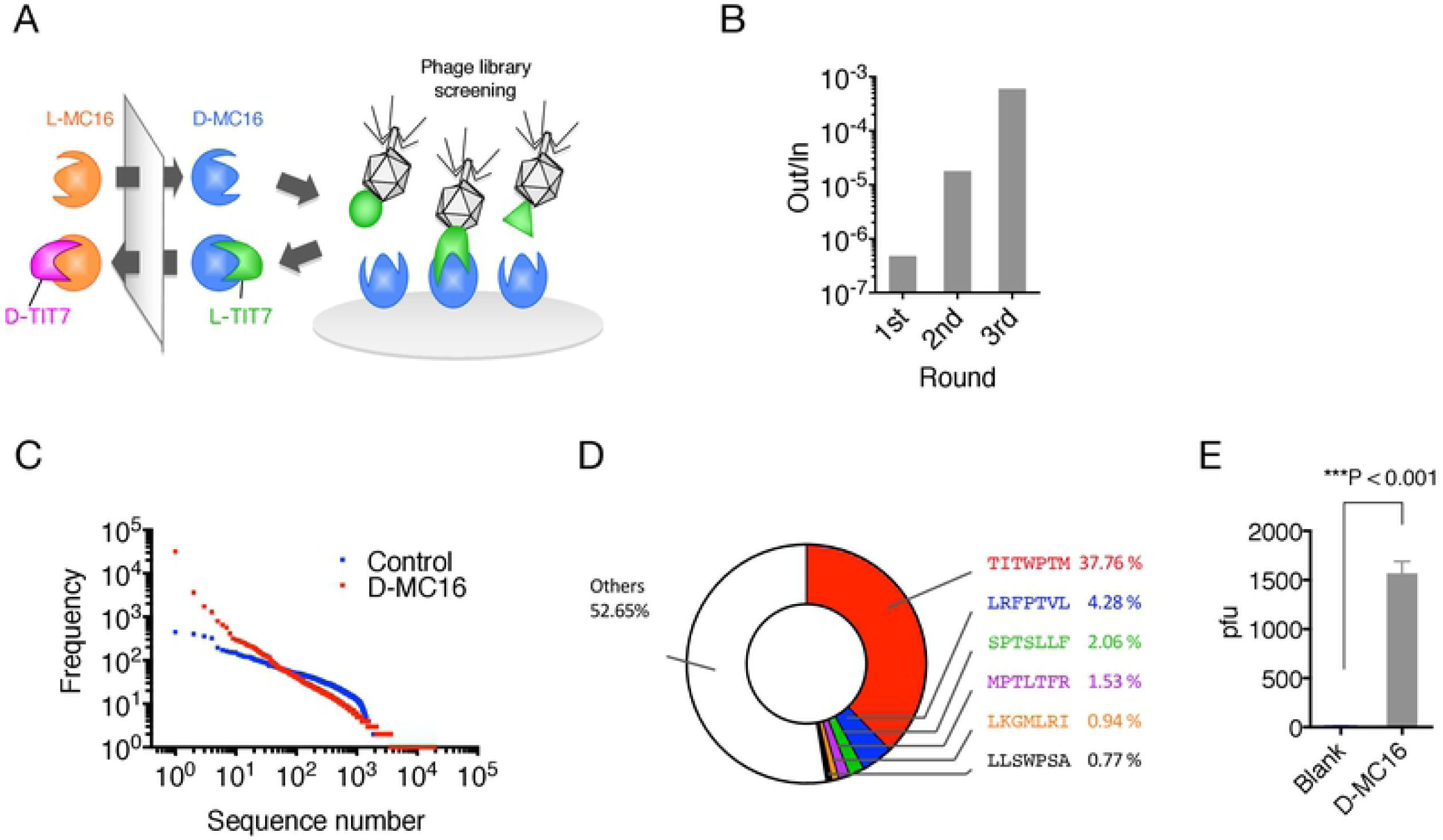
Mirror-image phage library screen for MC16-binding D-peptides. **A**. Strategy used to identify D-peptides using D-MC16 peptide as the target. **B**. Binding efficacy of phage pools obtained after each round, as assessed by plaque-forming assays. **C**. Proportion of peptides of various sequences in the third positive pool. The phage mixture was analyzed by next generation sequencing and ranked for peptide abundance (Supplemental Table 1). **D**. Distribution of peptide sequences in the third positive pool. **E**. Binding of phage clones displaying the TIT7 peptide sequence to D-MC16-versus control (blank)-coated plastic plates.

### Binding of dTIT7 to MC16 and ANXA1 *in vitro*

Since interaction of dTIT7 to ANXA1 N-terminal domain including MC16 likely occurs when MC16 is localized to the cell membrane, we mimicked this state by coating plastic plates with MC16 peptide and then adding a solution containing biotinylated dTIT7 to the plates. High levels of dTIT7 bound to WT MC16 in this context, with a Kd of 8.5 nM (Fig. 2A). We then assessed specificity of TIT7 binding to MC16 in a binding assay using mutant forms of MC16. That analysis indicated that dTIT7 binding affinity to MC16 mutants F7A, K9A and W11A was significantly lower than to WT MC16 (Fig. 2B).

**Fig. 2.**
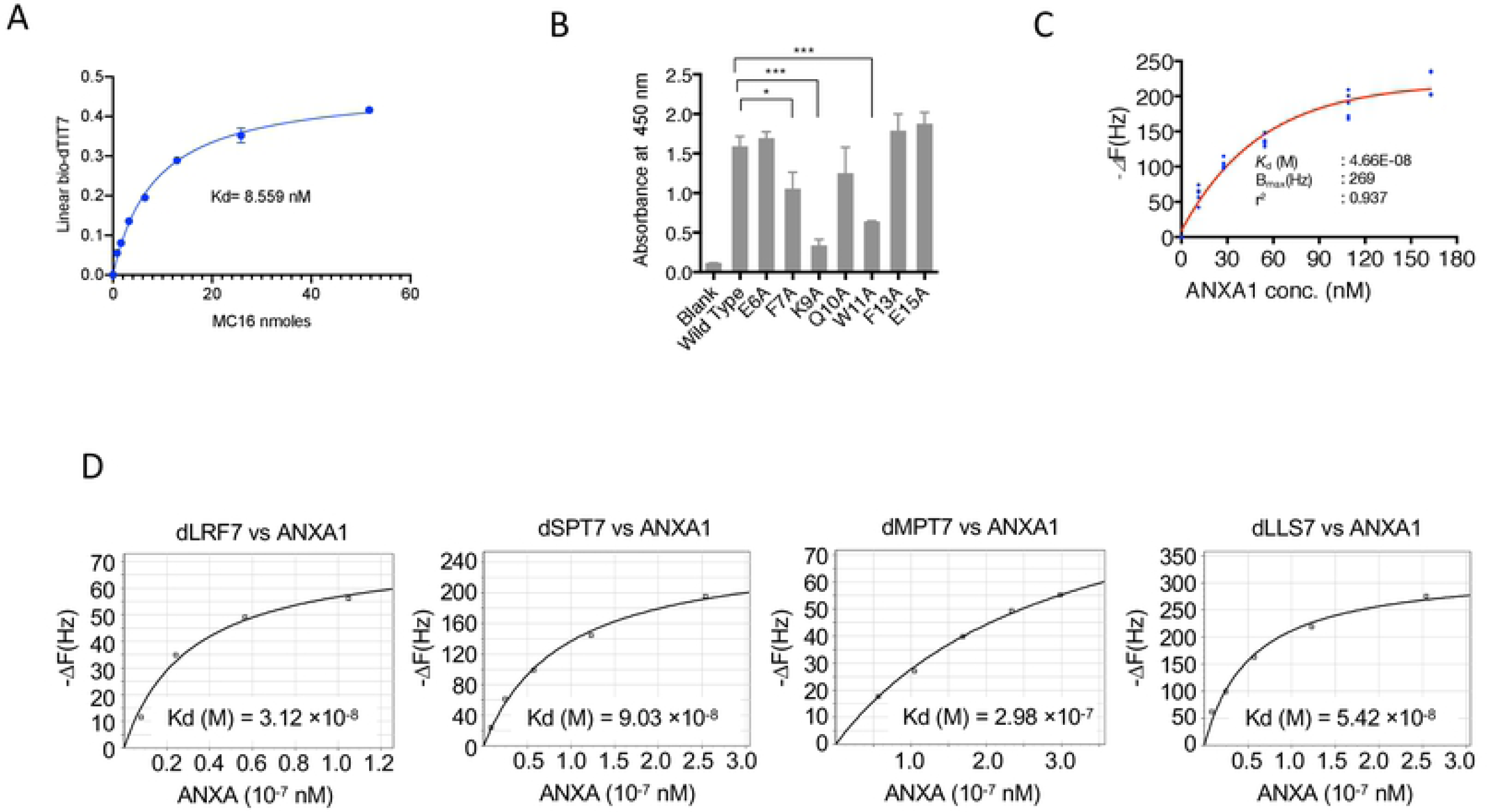
Binding of dTIT7 to an ANXA1 N-terminal domain peptide or to full-length ANXA1 protein. **A**. Plate binding assay of N-terminal biotinylated dTIT7 to synthetic human MC16 peptide, which represents the ANXA1 N-terminus. **B**. dTIT7 binding to human MC16 peptide and its mutants. In A and B, biotinylated dTIT7 peptide (1 μg/mL) was added to each MC16-coated plastic well and binding of peptide to MC16 was detected by a peroxidase-conjugated streptavidin and peroxidase color reaction. **C**. dTIT7 binding to recombinant full-length ANXA protein based on QCM analysis, which determines mass per unit area by measuring change in frequency of a dTIT7-coated sensor. **D**. Comparable QCM analysis relevant to other peptides identified in the screen.

We then assessed binding of full-length ANXA1 to immobilized dTIT7 by QCM analysis, which indicated a Kd of 4.66 × 10^−8^ M with ANXA1 (Fig. 2C), a value comparable to that for IF7 with ANXA1 (6.38 × 10^−8^M) [6]. QCM analysis of additional D-peptides identified in our screen (namely, d-LRF7, dSPT7, dMPT7 and dLLS7) with ANXA1 showed Kd values, ranging from 3-9 × 10^−8^ M (Fig. 2D), confirming that affinity of these peptides to ANXA1 was comparable to dTIT7 or IF7.

To confirm that dTIT7 and MC16 interact in solution, we analyzed a mixture of both peptides using NMR spectroscopy (Fig. 3A). We observed that the spectrum of the mixture was similar but differed in key ways from the sum of respective peptides. Most prominently, a distinct peak in the mixture spectrum emerged at 0.73 ppm (Fig. 3A, black arrow) and was absent in the summed spectrum. Concomitantly, we observed relative broadening of many peaks of the mixture spectrum. Note that a split in the peak at 1.08 ppm (Fig. 3A, red arrow) was relatively shallower in the mixture. Peak broadening has been attributed to shortened transverse relaxation time (*T*_2_) [13], which was indeed the case for the peak at 1.08 ppm (Fig. 3B). Moreover, *T*_2_ shortening is typically associated with an increase in molecular weight [13]. Overall, these results indicate an association between dTIT7 and MC16 in solution, at least at equilibrium.

**Fig. 3.**
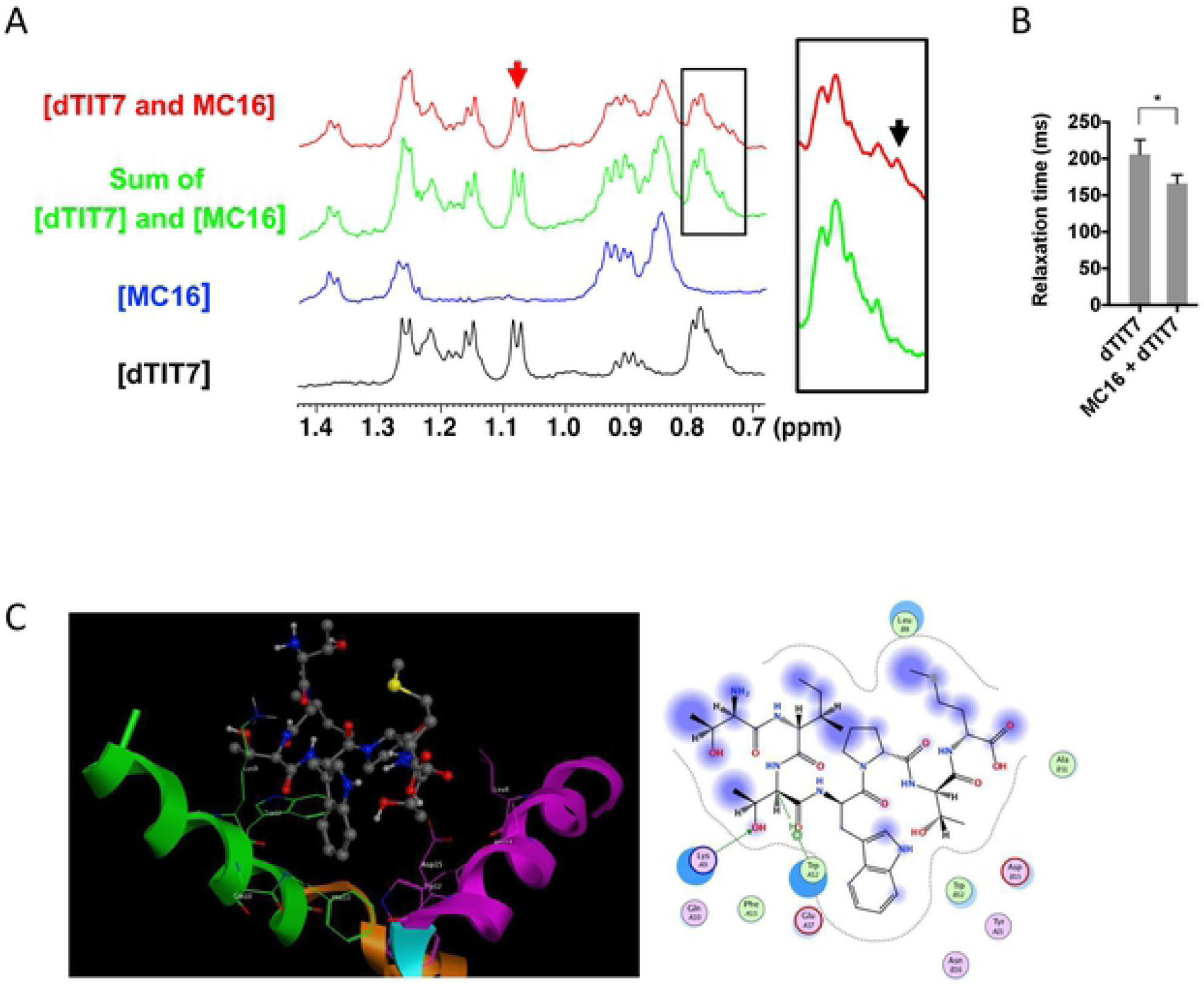
NMR analysis of dTIT7 interaction with monomeric L-MC16 in solution, and computer-simulated structure model of dTIT7 bound to the ANXA1 N-terminal domain. **A**. Shown are NMR spectra (methyl region) of dTIT7 (black line) or MC16 (blue line), the sum of both spectra (green line), and that of a mixture of both peptides (red line). Black arrow in square expanded at right indicates a peak at 0.73 ppm that emerged in the mixture spectrum, while red arrow indicates a peak at 1.08 ppm attributable to dTIT7. **B**. Transverse relaxation time (*T*_2_) of the dTIT7 peak at 1.08 ppm (red arrow in **c**) in the free state or in a mixture with MC16. **C**. Computer-simulated structural model for dTIT7 binding to the ANXA1 N-terminal domain. Proposed model was deduced by our previous study suggested that IF7 binds to an Anxa1 dimer [5], and results shown here suggest that MC16 polymerization is required for dTIT7 binding. The ANXA1 dimer structure was constructed by the Zdock module for protein-protein docking [9] and the 1HM6 X-ray structure of full-length ANXA1 was added to the 1MCX core domain at residue 40 [26, 27]. The modeled structure was then hydrogenated using the Protonate 3D module in MOE. After partial charges were assigned using the AMBER99 force field [28], hydrogen atoms were minimized. The dimer structure proposed here was ranked 37th in the top 2000 structures by this program. The Alpha Site Finder module in MOE was used to identify a potential IF7 binding pocket within the dimer. The proposed model was further validated by dG scoring calculated using MOE software with GBVI/WSA, a program allowing comparison of calculated and observed energetics [29]. The dTIT7 docking pose was calculated to be −5.1 kcal/mol.

We then generated a computer-simulated docking pose of dTIT7 with L-MC16 (Fig. 3C). To do so, we applied the strategy used to model IF7 binding with L-MC16 [6], in which two ANXA1 N-terminal domains provide a binding pocket for the ligand dTIT7. This model estimates the free energy of binding for dTIT7 to be −5.1 kcal/mol, while that for IF7 was estimated to be −3.7 kcal/mol [6].

### dTIT7 targeting of the brain tumor vasculature in mouse

We then used body imaging to confirm tumor vasculature-targeting activity of dTIT7 in brain tumor model mice using a conjugate of a near infra-red fluorescent reagent IRDye 800 CW to dTIT7 peptide. IRDye-dTIT7 was injected intravenously into brain-tumor bearing nude mice, and fluorescence was visualized using Xenogen IVIS imaging in real time, at various time points from 15 minutes to 144 hours (6 days) (Fig. 4A). IRdye-dTIT7 targeted brain tumor and kidney and remained detectable in these locations for up to 6 days after injection.

**Fig. 4.**
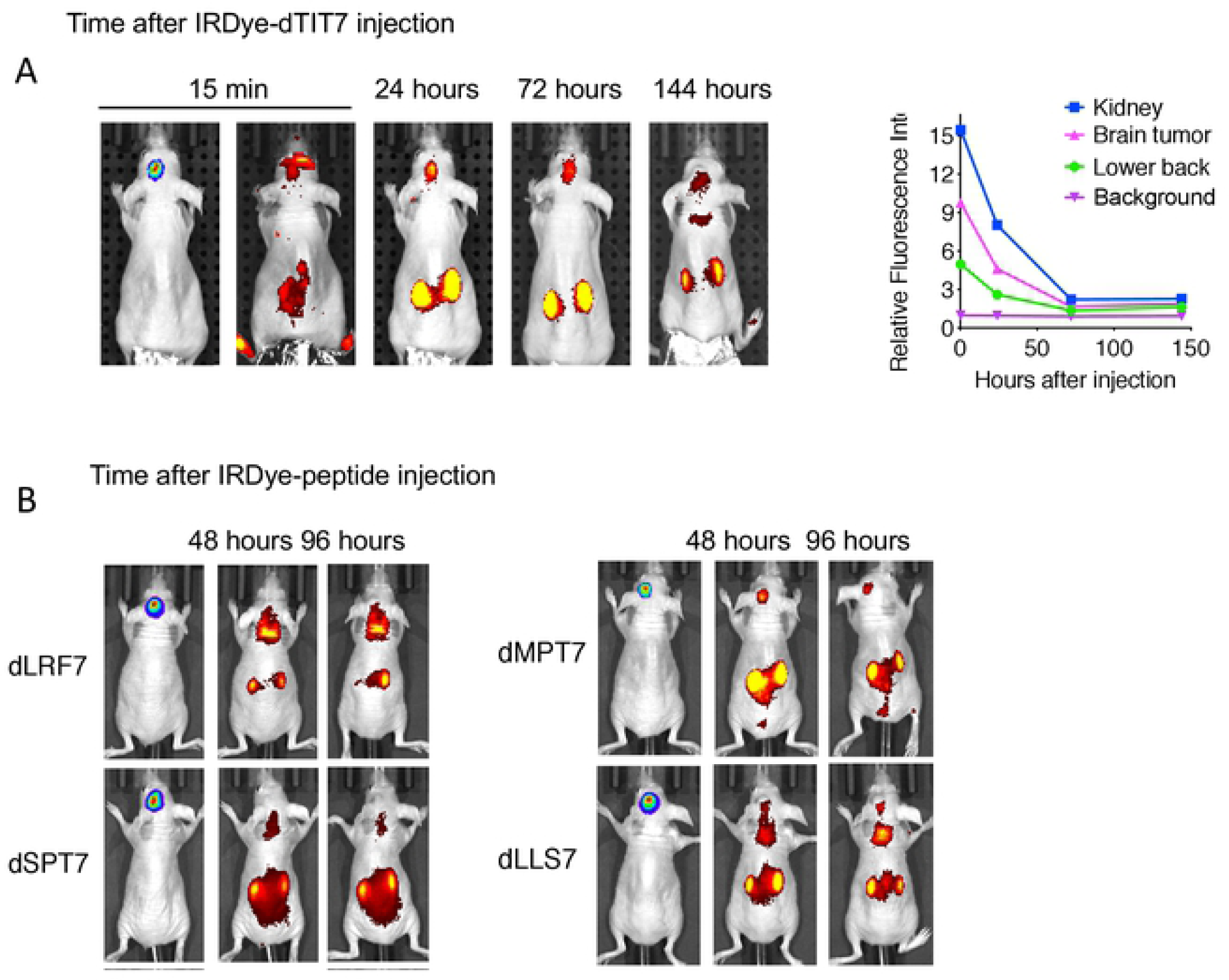
Whole body image analysis of IRDye-dTIT7 in brain tumor-bearing mice. **A**. Nude mice harboring C6-Luc brain tumors were injected with IRDye-dTIT7 through the tail vein. Whole body imaging of infra-red fluorescence was monitored using an IVIS imager. Right graph shows quantitative analysis of infra-red fluorescence signals in mice shown in left. **B**. Comparable whole body imaging for IRDye-conjugated dLRF7, dSPT7, dMPT7 and dLLS7.

In the same model, we also tested *in vivo* tumor vasculature-targeting of additional IRDye-conjugated peptides identified in our mirror-image phage library screen, namely d-LRF7, dSPT7, dMPT7 and dLLS7, using whole body imaging. That analysis revealed signals in brain, kidney and other organs (Fig. 4B). These results suggest that D-peptide sequences deduced in our screen targeted primarily the brain tumor and kidney vasculature.

### Therapeutic activity of dTIT7-conjugated GA

Previously, we conjugated IF7 with GA *via* non-cleavable linker [14]. Intravenously-injected GA-IF7 suppressed tumor growth in mouse breast, prostate, lung and melanoma tumor models [5]. Here, we prepared GA-dTIT7 as we had GA-IF7 [5] (Fig. 5) and determined its cytotoxic activity as well as that of control GA-C using C6 cells cultured *in vitro*. This assay showed the IC_50_ of GA-dTIT7 and GA-C to be 0.396 nM and 0.410 nM, respectively (Fig. 6A).

**Fig. 5.**
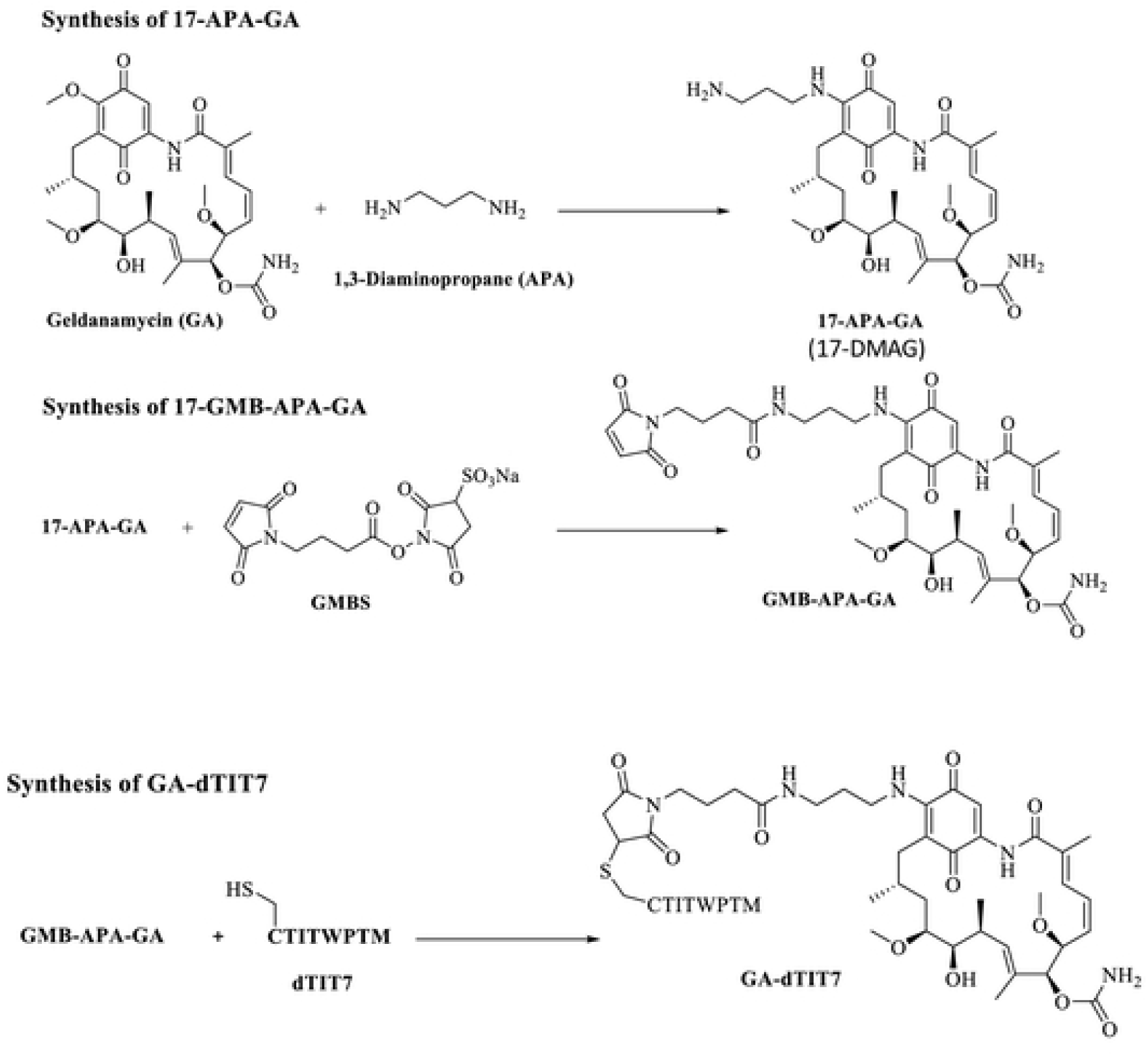
Three steps for the synthesis of GA-dTIT7. Procedures described by Mandler *et al*.[12] were modified as described in Materials and Methods. Note that 17-APA-GA is also known as 17-DMAG [15].

**Fig. 6.**
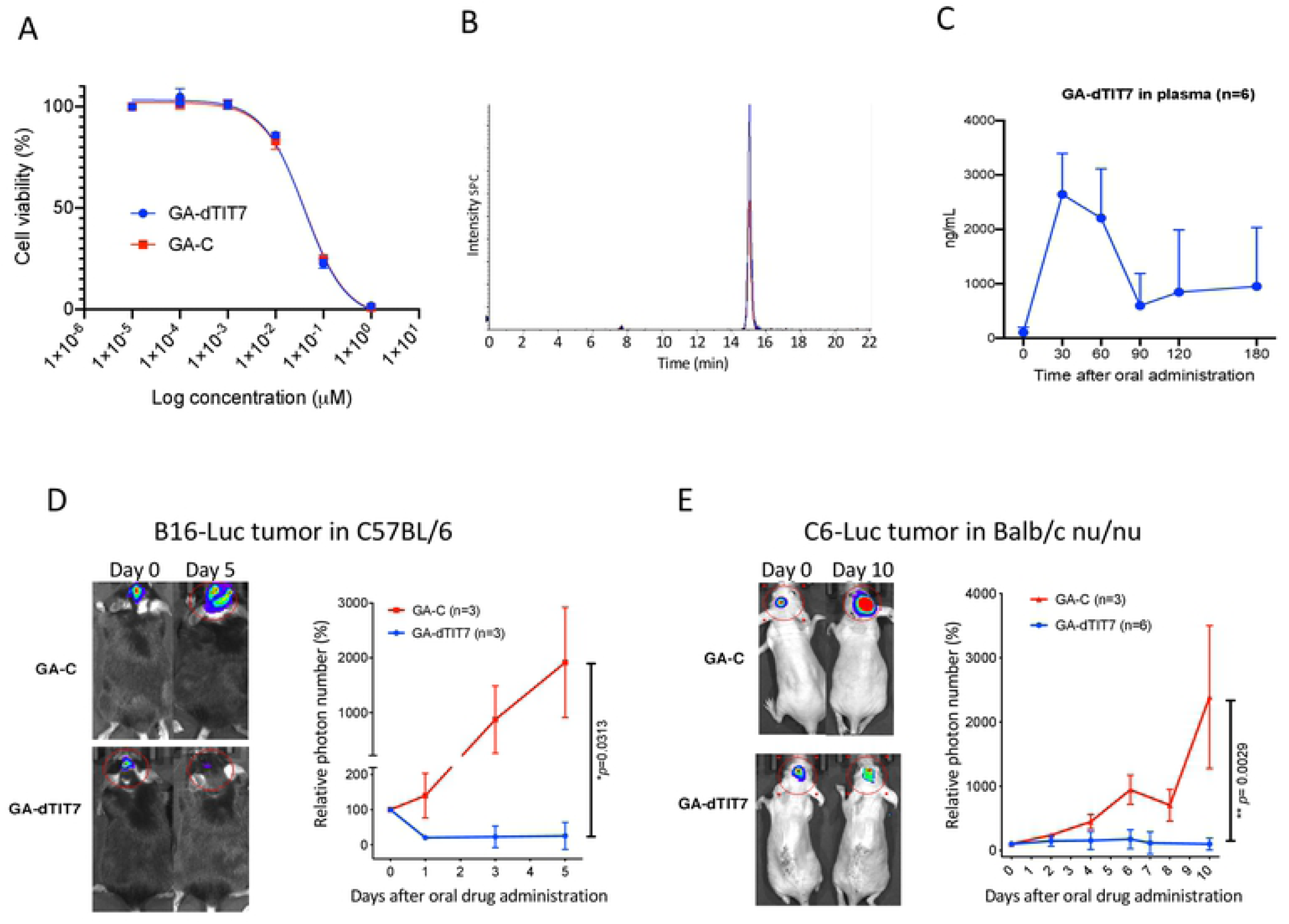
Therapeutic effect of orally administered GA-dTIT7 on brain tumors. **A**. Mouse melanoma B16 cells were treated with reagents shown at indicated concentrations, cultured 2 days, and assessed for viability using a CellTiter Glo (Promega) assay. The IC_50_ of each reagent was determined using GraphPad Prism program. B. Quantitative analysis of GA-dTIT7 in mouse plasma by LC-MS/MS. Plasma from GA-dTIT7-injected C57BL/6 female mice (9 μL) were combined with 1 μL GA-*iso*dTIT7 (1.0 μg), immediately mixed with 40 μL cold acetone, and then centrifuged to remove precipitates. The supernatant was then applied to LC-MS/MS, and eluates monitored by m/z 1725 for GA-*iso*dTIT7 (blue) and m/z 1719 for GA-dTIT7 (red). **C**. GA-dTIT7 levels in plasma from mice-orally administered GA-dTIT7. Each C57BL/6 mouse was orally-administered 1 mg GA-dTIT7. GA-dTIT7 levels were determined by LC-MS/MS, as shown in Supplemental Fig. 4. **C**. B16-Luc cells were injected into the brain of C57BL/6 mouse and tumor growth was monitored by IVIS imaging. When photon number reached 2 ×10^4 (approximately 5 days after B16-Luc cells inoculation), GA-dTIT7 (1.16 μmoles or 2 mg) or the molar equivalent GA-C (control) diluted with 10% taurodeoxycholate (200 μl) was orally-administered daily for 5 days. Panels at left show representative control and experimental mice imaged on days 0 and 5 after drug administration. Photon number is quantified at right. **D**. C6-Luc cells were injected into the brain of C57BL/6 mice and tumor growth was monitored by IVIS imaging. When photon number reached at 2 ×10^4 (approximately 10 days after C6-Luc cells inoculation), GA-dTIT7 and control GA-C were orally administered daily for 10 days. Panels at left show representative control and experimental mice imaged on days 0 and 10 after drug administration. Photon number is quantified at right. In these graphs, error bars denote means ± SEM. Statistical analysis was assessed by Student’s t-test.

In our previous study we found that intravenously-injected GA-IF7 at 6.5 μmoles/kg suppressed growth of melanoma, lung carcinoma, prostate cancer, and breast cancer models in the mouse [5]. When we injected GA-dTIT7 at 6.5 μmoles/kg intravenously to the tumor-bearing mice in the same manner as we have done for GA-IF7, GA-dTIT7 did not suppress tumor growth (data not shown). Since it is known that the GA analogue 17-DMAG, which is a part of GA-dTIT7 (Fig. 5), is orally-administrable [15], we asked if orally-administered GA-dTIT7 enters the circulation by assessing gut-to-blood GA-dTIT7 transport using quantitative LC-MS/MS analysis of isotopically-labeled dTIT7 (*iso*dTIT7). The molecular weight of GA-*iso*dTIT7, in which the last methionine residue contains ^13^C and ^15^N, is 1725.60 Da, while that of the internal standard, GA-dTIT7, is 1719.6 Da (Fig. 6B). For this analysis, we dissolved GA-dTIT7 in 10% taurodeoxycholate (TDC) in water to enhance drug transport from the digestive tract to the circulation more efficiently than GA-dTIT7 formulated with 10% Solutol HS15, 6% glucose or 10% carboxymethyl cellulose. Plasma samples from mice orally-administered GA-dTIT7 were combined with GA-*iso*dTIT7.Then after removal of proteins by precipitation with cold acetone, we subjected the supernatant to LC-MS/MS analysis to determine the quantity of GA-dTIT7. This analysis showed a time-dependent increase in GA-dTIT7 in mouse plasma, peaking at 30 min (Fig. 6C). When 1 mg GA-dTIT7 was orally-administered, the plasma concentration drug at 30 min was 2.62 ± 0.69 ng /mL, or 1.52 nM.

Next, we tested the therapeutic effect of orally-administered GA-dTIT7 on brain tumors *in vivo*. We had previously shown that IF7-SN38 overcame the BBB and suppressed brain tumor growth in model mice [6]. To determine whether GA-dTIT7 functioned similarly, we established B16-Luc tumors in brains of C57BL/6 mice and monitored tumor growth by photon number produced by luciferase using IVIS imaging. When photon number reached 1×10^4, we orally administered GA-dTIT7 (1 mg or 0.58 μmoles) in 200 μL in 10% TDC in water daily for 7 days but did not observe suppression of tumor growth (data not shown). However, when we doubled the GA-dTIT7 dose to 2 mg and orally-administered the drug daily for 5 days, imaging revealed significant suppression of tumor growth in GA-dTIT7-treated mice, while tumors continued to grow in control mice that had received 1 mg GA-C (the molar equivalent of GA-dTIT7) daily for 5 days (Fig. 6D). Comparable analysis using C6-Luc brain tumor models in nude mice revealed tumor growth suppression by GA-dTIT7 but not control GA-C (Fig. 6E). These results showed, as a proof-of-concept, that orally-administered GA-dTIT7 suppresses brain tumors *in vivo* in mice.

## Discussion

Here we used a mirror-image peptide display strategy [8] to identify a series of linear 7-mer D-peptides using the ANXA1 NH_2_-terminal domain peptide (Fig. 1A). Because this strategy requires a chemically synthesized receptor made of D-amino acids, the application is limited to proteins in which a synthetic version of the peptide functions as receptor for the protein of interest. Nonetheless, this strategy have been successfully applied to develop therapeutic D-peptide modulators of the tyrosine kinase SH3 domain [7] or inhibitors of amyloid beta aggregation in Alzheimer’s disease [16]. In both cases, each D-target conformed to a unique stereo-specific structure and provided a binding pocket for L-peptides displayed on the phage. In our study, we also exploited the fact that a chemically-synthesized peptide representing the ANXA1 NH_2_-terminal domain (MC16) served as receptor for IF7 [6]. Although MC16 is considered too short and flexible in solution to form a stable 3-D structure, IF7/MC16 interactions were detected in our binding assays, including a plate binding assay, fluorescence correlation spectroscopy, and QCM [6]. Indeed, D-peptides identified by a D-MC16 target also bound to L-MC16 and full-length ANXA1 protein (Figs. 2 and 3).

Others have reported a D-peptide alternative for IF7 designated retro-inverso IF7 (RIF7), in which the reverse IF7 sequence was synthesized using D-amino acids [17]. When RIF7 was conjugated to red fluorescent 5-carboxytetramethylrhodamine (TMR) and then injected intravenously into a pulmonary cancer model mouse, TMR-RIF7 targeted the lung tumor and exhibited prolonged stability compared to TMR-IF7 [17]. That study showed that TMR-IF7 and TMR-RIF7 targeted not only tumors but also several normal organs. Such non-specific organ targeting is likely due partially to TMR, as green fluorescent Alexa 488-labeled IF7 targeted brain tumors but not to the normal organs [6]. Thus far no one has reported a therapeutic effect of a RIF7-conjugated drug.

Currently, tumors are often diagnosed by positron emission tomography (PET) scans utilizing radioactive ^18^F glucose or FDG. Despite the highly specific tumor vasculature targeting activity of IF7, our attempts to conduct PET with IF7 were not successful (data not shown), although others have shown detectable, though limited, tumor imaging with IF7 [18-20]. We emphasize, however, that whole body imaging of IRDye-conjugated dTIT7 indicated clear brain tumor targeting (Fig. 4). Compared with other organ systems, FDG-PET imaging of the brain presents unique challenges because of high background glucose metabolism in normal gray matter [21]. We consider that D-peptides identified here warrant further testing in imaging of brain tumors.

Although we had anticipated that intravenously-injected GA-dTIT7 would exhibit anti-tumor activity *in vivo*, we did not observe therapeutic activity of dTIT7-conjugated drugs following intravenous injection, suggesting that either higher dosages of GA-dTIT7 or different drug formulation may be required. Relevant to the latter, detergents significantly alter IF7-SN38 therapeutic efficacy: we have shown that formulation with 10% Solutol in water significantly reduces the effective dosage against brain tumors [6]. Future studies should address these issues in the case of GA-dTIT7 following intravenous injection.

GA analogues 17-AAG and 17-DMAG have been shown to be potent anti-cancer agents with less toxicity than the parental drug GA. However, several clinical trials with these GA analogues indicated toxicity too high to proceed beyond a phase II trial [22, 23]. Nonetheless, pre-clinical and clinical studies of the GA analogue 17-DMAG showed it is orally-administrable [15, 24]. We found that GA-dTIT7 (Fig. 5) is orally administrable and suppressed tumor growth in mouse brain tumor models (Fig. 6 DE). We were able to test oral administration of GA-dTIT7 as this compound exhibits cytotoxic activity (Fig 6A). GA-dTIT7 should be stable *in vivo*, as GA is linked to dTIT7 through an esterase-resistant linker and dTIT7 is expected to be resistant to digestive proteases. Although the efficacy of GA-dTIT7 gut-to-blood transport was low here (Fig. 6C), future studies should address how to improve this efficacy. Additional modification of GA-dTIT7 to enhance ANXA1-binding and gut-to-blood transfer activities could strengthen the clinical relevance of this drug.

Cancer treatments are increasingly expensive due to development of sophisticated diagnostics and therapies. Our drug, which consists of a short peptide plus an anti-cancer reagent, can be chemically synthesized cost-effectively. Given that ANXA1 is an extremely specific tumor vasculature surface marker [2], and IF7-conjugated anti-cancer drugs have profound effects on subcutaneous and brain tumors [5, 6, 25], a drug conjugated to an ANXA1-binding peptide should eradicate tumors effectively at low dosage and minimize side effects. Finally, orally-administrable drugs would be advantageous in economically disadvantaged societies that lack infrastructure required for costly treatment. As clinical trials with tumor vasculature-homing peptides are beginning, we will soon be able to evaluate efficacy of these strategies in cancer patients. Further development of peptide-conjugated drugs could reveal strong candidates for clinical applications to treat intractable cancers.

## Acknowledgement

This study was supported by an institutional grant LEAD at National Institute of Advanced Industrial Science and Technology, by the Project for Cancer Research and Therapeutic Evolution (P-CREATE) from Japan Agency for Medical Research and Development (AMED) to MNF, by P41 GM103390 and P41 RR005351. NM is a recipient of a Research Grant for Young Japanese Scientists from The Nakajima Foundation. We thank Dr. Elise Lamar for editing the manuscript and Mrs. Hisae Okuhara for clerical/administrative assistance.

## Supporting Information

**S1 Table. Nucleotide and peptide sequences obtained by NGS (next generation sequencing) of the third positive phage pool**.

## References

1. Ruoslahti E, Rajotte D. An address system in the vasculature of normal tissues and tumors. Annu Rev Immunol. 2000;18:813–27. PubMed PMID: 10837076.

2. Oh P, Li Y, Yu J, Durr E, Krasinska KM, Carver LA, et al. Subtractive proteomic mapping of the endothelial surface in lung and solid tumours for tissue-specific therapy. Nature. 2004;429(6992):629–35. PubMed PMID: 15190345.

3. Oh P, Testa JE, Borgstrom P, Witkiewicz H, Li Y, Schnitzer JE. In vivo proteomic imaging analysis of caveolae reveals pumping system to penetrate solid tumors. Nat Med. 2014;20(9):1062–8. doi: 10.1038/nm.3623. PubMed PMID: 25129480.

4. Hatakeyama S, Sugihara K, Nakayama J, Akama TO, Wong SM, Kawashima H, et al. Identification of mRNA splicing factors as the endothelial receptor for carbohydrate-dependent lung colonization of cancer cells. Proc Natl Acad Sci U S A. 2009;106(9):3095–100. PubMed PMID: 19218444; PubMed Central PMCID: PMC264266.

5. Hatakeyama S, Sugihara K, Shibata TK, Nakayama J, Akama TO, Tamura N, et al. Targeted drug delivery to tumor vasculature by a carbohydrate mimetic peptide. Proc Natl Acad Sci U S A. 2011;108(49):19587–92. Epub 2011/11/25. doi: 1105057108 [pii] 10.1073/pnas.1105057108. PubMed PMID: 22114188; PubMed Central PMCID: PMC3241764.

6. Nonaka M, Suzuki-Anekoji M, Nakayama J, Mabashi-Asazuma H, Jarvis DL, Yeh JC, et al. Overcoming the blood-brain barrier by Annexin A1-binding peptide to target brain tumours. Br J Cancer. 2020. Epub 2020/09/15. doi: 10.1038/s41416-020-01066-2. PubMed PMID: 32921792.

7. Schumacher TN, Mayr LM, Minor DL, Jr., Milhollen MA, Burgess MW, Kim PS. Identification of D-peptide ligands through mirror-image phage display. Science. 1996;271(5257):1854–7. Epub 1996/03/29. PubMed PMID: 8596952.

8. Funke SA, Willbold D. Mirror image phage display--a method to generate D-peptide ligands for use in diagnostic or therapeutical applications. Mol Biosyst. 2009;5(8):783–6. doi: 10.1039/b904138a. PubMed PMID: 19603110.

9. Chen R, Li L, Weng Z. ZDOCK: an initial-stage protein-docking algorithm. Proteins. 2003;52(1):80–7. Epub 2003/06/05. doi: 10.1002/prot.10389. PubMed PMID: 12784371.

10. Jarvis DL. Baculovirus-insect cell expression systems. Methods Enzymol. 2009;463:191–222. Epub 2009/11/07. doi: 10.1016/S0076-6879(09)63014-7. PubMed PMID: 19892174.

11. Suzuki M, Nakayama J, Suzuki A, Angata K, Chen S, Sakai K, et al. Polysialic acid facilitates tumor invasion by glioma cells. Glycobiology. 2005;15(9):887–94. PubMed PMID: 15872150.

12. Mandler R, Dadachova E, Brechbiel JK, Waldmann TA, Brechbiel MW.Synthesis and evaluation of antiproliferative activity of a geldanamycin-Herceptin immunoconjugate. Bioorg Med Chem Lett. 2000;10(10):1025–8. PubMed PMID: 10843208.

13. Wüthrich K. NMR of Proteins and Nucleic Acids. New York: John Wiley & Sons; 1986.

14. DeBoer C, Meulman PA, Wnuk RJ, Peterson DH. Geldanamycin, a new antibiotic. J Antibiot (Tokyo). 1970;23(9):442-7. Epub 1970/09/01. doi: 10.7164/antibiotics.23.442. PubMed PMID: 5459626.

15. Ikebe E, Kawaguchi A, Tezuka K, Taguchi S, Hirose S, Matsumoto T, et al. Oral administration of an HSP90 inhibitor, 17-DMAG, intervenes tumor-cell infiltration into multiple organs and improves survival period for ATL model mice. Blood Cancer J. 2013;3:e132. Epub 2013/08/21. doi: 10.1038/bcj.2013.30. PubMed PMID: 23955587; PubMed Central PMCID: PMCPMC3763384.

16. Wiesehan K, Buder K, Linke RP, Patt S, Stoldt M, Unger E, et al. Selection of D-amino-acid peptides that bind to Alzheimer’s disease amyloid peptide abeta1-42 by mirror image phage display. Chembiochem. 2003;4(8):748–53. Epub 2003/08/05. doi: 10.1002/cbic.200300631. PubMed PMID: 12898626.

17. Chen X, Fan Z, Chen Y, Fang X, Sha X. Retro-inverso carbohydrate mimetic peptides with annexin1-binding selectivity, are stable in vivo, and target tumor vasculature. PLoS One. 2013;8(12):e80390. doi: 10.1371/journal.pone.0080390. PubMed PMID: 24312470; PubMed Central PMCID: PMC3846562.

18. Chen F, Pu X, Xiao Y, Shao K, Wang J, Hu W, et al. Preparation and SPECT imaging of the novel Anxa 1-targeted probe 99mTc-p-SCN-Bn-DTPA-GGGRDN-IF7. Journal of Radioanalytical and Nuclear Chemistry. 2019;320:525–30. doi: doi.org/10.1007/s10967-019-06500-1.

19. Chen F, Xiao Y, Shao K, Zhu B, Jianng M. Positron emission tomography imaging of a novel Anxa1-targeted peptide 18F-Al-NODA-Bn-p-SCN-GGGRDNIF7 in A431 cancer mouse models. J Label Compd Radiopharm. 2020:1–8. doi: 10.1002/jlcr.3865.

20. Gu X, Jiang M, Pan D, Cai G, Zhang R, Zhou Y, et al. Preliminary evaluation of novel 18F-AlF-NOTA-IF7 as a tumor imaging agent. Radioanal Nucl Chem. 2016;308:851–6. doi: 10.1007/s10967-015-4533-3.

21. Wong TZ, van der Westhuizen GJ, Coleman RE. Positron emission tomography imaging of brain tumors. Neuroimaging Clin N Am. 2002;12(4):615–26. Epub 2003/04/12. PubMed PMID: 12687915.

22. Ronnen EA, Kondagunta GV, Ishill N, Sweeney SM, Deluca JK, Schwartz L, et al. A phase II trial of 17-(Allylamino)-17-demethoxygeldanamycin in patients with papillary and clear cell renal cell carcinoma. Invest New Drugs. 2006;24(6):543–6. PubMed PMID: 16832603.

23. Kang MH, Reynolds CP, Houghton PJ, Alexander D, Morton CL, Kolb EA, et al. Initial testing (Stage 1) of AT13387, an HSP90 inhibitor, by the pediatric preclinical testing program. Pediatr Blood Cancer. 2012;59(1):185–8. doi: 10.1002/pbc.23154. PubMed PMID: 21538821; PubMed Central PMCID: PMCPMC3154460.

24. Li YP, Gao LY, Li KT, Meng S, Zhu JH, Li D, et al. LC-MS/MS method for determination of geldanamycin derivative GM-AMPL in rat plasma to support preclinical development. J Chromatogr B Analyt Technol Biomed Life Sci. 2013;912:43–9. Epub 2012/12/25. doi: 10.1016/j.jchromb.2012.09.002. PubMed PMID: 23261821.

25. Yu DH, Liu YR, Luan X, Liu HJ, Gao YG, Wu H, et al. IF7-Conjugated Nanoparticles Target Annexin 1 of Tumor Vasculature against P-gp Mediated Multidrug Resistance. Bioconjug Chem. 2015;26(8):1702–12. Epub 2015/06/16. doi: 10.1021/acs.bioconjchem.5b00283. PubMed PMID: 26076081.

26. Shesham RD, Bartolotti LJ, Li Y. Molecular dynamics simulation studies on Ca2+ - induced conformational changes of annexin I. Protein Eng Des Sel. 2008;21(2):115–20. Epub 2008/02/20. doi: 21/2/115 [pii] 10.1093/protein/gzm094. PubMed PMID: 18283055.

27. Rosengarth A, Gerke V, Luecke H. X-ray structure of full-length annexin 1 and implications for membrane aggregation. J Mol Biol. 2001;306(3):489–98. Epub 2001/02/17. doi: 10.1006/jmbi.2000.4423 S0022-2836(00)94423-1 [pii]. PubMed PMID: 11178908.

28. Kollman PA, Massova I, Reyes C, Kuhn B, Huo S, Chong L, et al. Calculating structures and free energies of complex molecules: combining molecular mechanics and continuum models. Acc Chem Res. 2000;33(12):889–97. Epub 2000/12/22. doi: ar000033j [pii]. PubMed PMID: 11123888.

29. Corbeil CR, Williams CI, Labute P. Variability in docking success rates due to dataset preparation. J Comput Aided Mol Des. 2012;26(6):775–86. Epub 2012/05/09. doi: 10.1007/s10822-012-9570-1. PubMed PMID: 22566074; PubMed Central PMCID: PMC3397132.

